# Monitoring conservation effectiveness in tropical forests: bushmeat offtake parameters vary more quickly compared to parameters related to the living wildlife community

**DOI:** 10.1101/2022.02.04.479098

**Authors:** Jacques Keumo Kuenbou, Nikki Tagg, Donald Mbohli Khan, Stjin Speelman, Jacob Willie

## Abstract

Evaluating the effectiveness of conservation actions in tropical forests is essential. Generally based on the monitoring of living wildlife communities, combination with bushmeat extraction indicators is seldom used. It is therefore interesting to carry out a monitoring of indices related to these two categories of indicators in order to identify indices presenting more temporal variation and allowing for a rapid detection of real changes. Between 2017 and 2020, data on bushmeat offtake were recorded and surveys of wildlife and human activity signs were conducted in a conservation zone in Dja Biosphere Reserve in Cameroon. Data were collected around villages where human pressure is high. Our results show a decline in total biomass and number of animals killed. An increase in numbers of traps set was observed, but there was no notable difference in the mean biomass of a carcass and mean number of hunting trips. Overall, wildlife abundance decreased slightly in 2020, mainly reflecting changes for three species—bay duiker (Cephalophus dorsalis), yellow-backed duiker (Cephalophus sylvicultor) and red river hog (Potamocherus porcus)—which were more abundant in 2018. Regarding keystone species, we noted a slight increase in the abundance of chimpanzees. Patterns of species richness in both study years were similar. These results suggest that bushmeat offtake data may be more informative for early evaluations of the effectiveness of wildlife conservation efforts. This underlines the importance of monitoring bushmeat for the evaluation of wildlife conservation projects in contexts where hunting occurs.

## Introduction

In tropical forests around the world, hunting is recognized as one of the greatest threats to biodiversity (Leverington et al. 2010; Abernethy et al. 2013; Maxwell et al. 2016). This is especially the case for African forests (Fa et al. 2002; Abernethy et al. 2013). Hunting for wild meat known as “bushmeat” is a major threat in Central Africa and harvest estimates in that area range from one (Wilkie and Carpenter 1999) to five million tons per year (Fa et al. 2003). For centuries, bushmeat has been used as a food source for African populations (Elliott et al. 2002; Milner-Gulland and Bennett 2003; Fa and Brown 2009) and remains the principal source of protein (Wilkie et al. 2005; Sirén and Machoa 2008; Nasi et al. 2011; Barboza et al. 2016). In rural areas, it was estimated that people obtain at least 20% of their protein from wild animals (Chardonnet et al. 1995). The livelihoods of rural inhabitants also depend upon these resources as significant source of revenue (Brown 2003; Brown and Williams 2003; Milner-Gulland and Bennett 2003; De Merode et al. 2004). Hunting in these areas is an important commercial activity that can provide substantial benefits in a context with few alternative cash-generating opportunities (E. L. Bennett and J. G. Robinson 2000; Fa 2007). Rapid human population growth, socioeconomic change, infrastructure development, technological improvements and rural poverty in African countries create increased pressure on wildlife and other natural resources (Bennett et al. 2007; Barrett et al. 2011; Abernethy et al. 2013; Van Zyl 2015; Lindsey et al. 2017). This pressure is accentuated by the growing demand from urban populations who, despite having alternative sources of protein, sometimes consider bushmeat to be a luxury good (van Vliet and Mbazza 2011). This pressure could result in long-term modifications to forest ecosystem dynamics— through the reduction of seed dispersers like granivores and frugivores which play a crucial role in the recruitment of large-seeded plants (Vanthomme et al. 2010)—and therefore knock-on effects for global climate regulation (Devaraju et al. 2015) and climate change (Lewis 2006; Lewis et al. 2015).

Threats to biodiversity underline the necessity to undertake conservation actions. It is now widely recognized that poverty in rural people living around protected areas could be a brake to conservation (Adams et al. 2004). The Socio-economic development of people living around protected areas is highly linked to wildlife trends and is critical for the successful maintenance of wildlife populations (Barnes et al. 2016). In order to reduce the impact on species, conservation projects should therefore take into account local people in wildlife management by providing alternatives to threats caused by anthropogenic activity. In Africa in particular, a major threat is hunting. The development of mechanisms to increase rural people’s income should be an integral part of conservation projects. Several options can be considered regarding activities to be settled, direct payment to communities for environmental services (Ferraro 2001; Raes 2016) and indirect payment through support in the development of income-generating activities such as community-based ecotourism (Gubbi et al. 2008; Lovett and Okwell 2010) and support in agricultural production.

Evaluating the effectiveness of such community-level conservation efforts is important in order to identify if actions are having a positive impact on wildlife conservation (Sebastián-González et al. 2011; Bottrill and Pressey 2012). Tools have been developed in order to define the evaluation criteria of projects according to the threats and actions conducted (Salafsky et al. 2002; Salafsky et al. 2008), helping to better evaluate the causal links between action and outcome (Chaves et al. 2018; Qiu et al. 2018). However, there is a lack of knowledge on the effectiveness of different conservation actions, and of monitoring and evaluation (Sutherland et al. 2004; Ferraro and Pattanayak 2006), particularly in Central African forests. Nevertheless, some conservation projects have been evaluated through monitoring of species population dynamics (Williams et al. 1999). Other evaluation methods involve the analysis of the cost-effectiveness of conservation actions (Sebastián-González et al. 2011; Raes et al. 2014) and the evaluation of income generated by local people in projects (Raes et al. 2016). However, these methods do not evaluate the impact on the ecosystems to be protected. Another way to evaluate projects is to interview project managers (Wicander and Coad 2018), but this approach does not take into account practical evidence (Sutherland et al. 2004). For many years, evaluating the impact of these conservation actions by managers and researchers was simply done through the monitoring of trends and status of the biodiversity they were trying to conserve (Salafsky et al. 2002). However, measuring biodiversity status of (species, habitats, ecosystems) alone often fails to demonstrate any impact of these actions (Salzer and Salafsky 2006).

There are no systematic indicators to measure success of conservation projects. Although not sufficient to fully evaluate community-level conservation efforts, ecological monitoring of species populations is essential for long-term conservation project evaluation (Kremen et al. 1994). Wildlife population change is an important and useful metric for evaluating wildlife conservation outcomes in and outside protected areas. It is sensitive to long-term environmental change (Buckland et al. 2005), often directly linked to conservation objectives, and valuable in diagnosing extinction risk (Hoffmann et al. 2015). Several methods of biodiversity measurement have been developed, making it possible to assess the impact of natural and anthropogenic factors on biodiversity (Cury and Christensen 2005; Shin and Shannon 2010). These methods range from estimating relative abundances, which is less precise with regard to determining the role of species in the ecosystem (Hoffmann et al. 2015), to calculating the various functional diversity indices which takes into account how environmental disturbances affect biodiversity and what are the consequences for ecosystem functioning (Petchey and Gaston 2006; Camara 2016; Oliveira et al. 2016; Mazel et al. 2018). Bushmeat is a crucial issue for both wildlife conservation and human development, and initiatives to tackle the crisis need to provide for both of these, often opposing needs (Coad et al. 2007). Hence, evaluation of the effectiveness of conservation projects in areas where hunting is the main threat must be done through the monitoring of wildlife harvesting. We therefore have two main groups of indicators, the first related to the living wildlife community in the forest (abundance, species richness and functional diversity) and the second consisting of bushmeat extraction data. Conservation effort evaluations are generally based on one of two groups of indices (often only on wildlife abundance), and conservation projects to be evaluated are usually implemented for relatively short periods. It is therefore relevant to monitor these indices in order to determine how they covary and identify indices that present more temporal variations and allow for a rapid detection of real changes in the short term.

The overall goal of this research, therefore, is to assess how bushmeat offtake and wildlife community structure covary through time. Specifically, we will evaluate trends in bushmeat harvesting dynamics over time, determine trends in wildlife abundance, species richness, and composition over the years. We predict that the number of species hunted, number of carcasses, amount of harvested biomass, and hunting effort will display a more rapid change than wildlife abundance, species richness, and functional diversity which will remain constant over years.

## Materials and methods

### Study area

The study was carried out in the northern periphery of the Dja Biosphere Reserve (DBR) in the eastern region of Cameroon (Fig 1). This area is located in lowland rainforest where there are two rainy seasons and two dry seasons. Recent climatic data show an annual precipitation average 1637.9 mm (SD=105.1 mm) and an average temperature around 23°C (Willie et al. 2014). Within the target area are 17 villages running along the edge of the Dja River and harboring around 250 households. A project was implemented here for 4 years (March 2017–March 2021) with the aim to enable rural poor to help protect biodiversity. The vision of the project was to reduce hunting pressure in the area by supporting local communities in the development of alternative income-generating activities, because local people in the area hunt mainly for sale in order to obtain money because they do not have other income sources (Epanda et al. 2019). Agreements were signed between individual villagers and some conservation organizations operating in the area (“Tropical Forest and Rural Development”, “Fondation Camerounaise Terre Vivante”. In these agreements, the individual villager commits to reducing its hunting activities in exchange for support and accompaniment from these organizations and development of other income-generating activities. This support is provided through technical and financial backing for the creation and extension of cocoa plantations as well as the supply of fishing equipment. Traditionally, cocoa and fishing have been important sources of income in this area.

**Figure 1:**
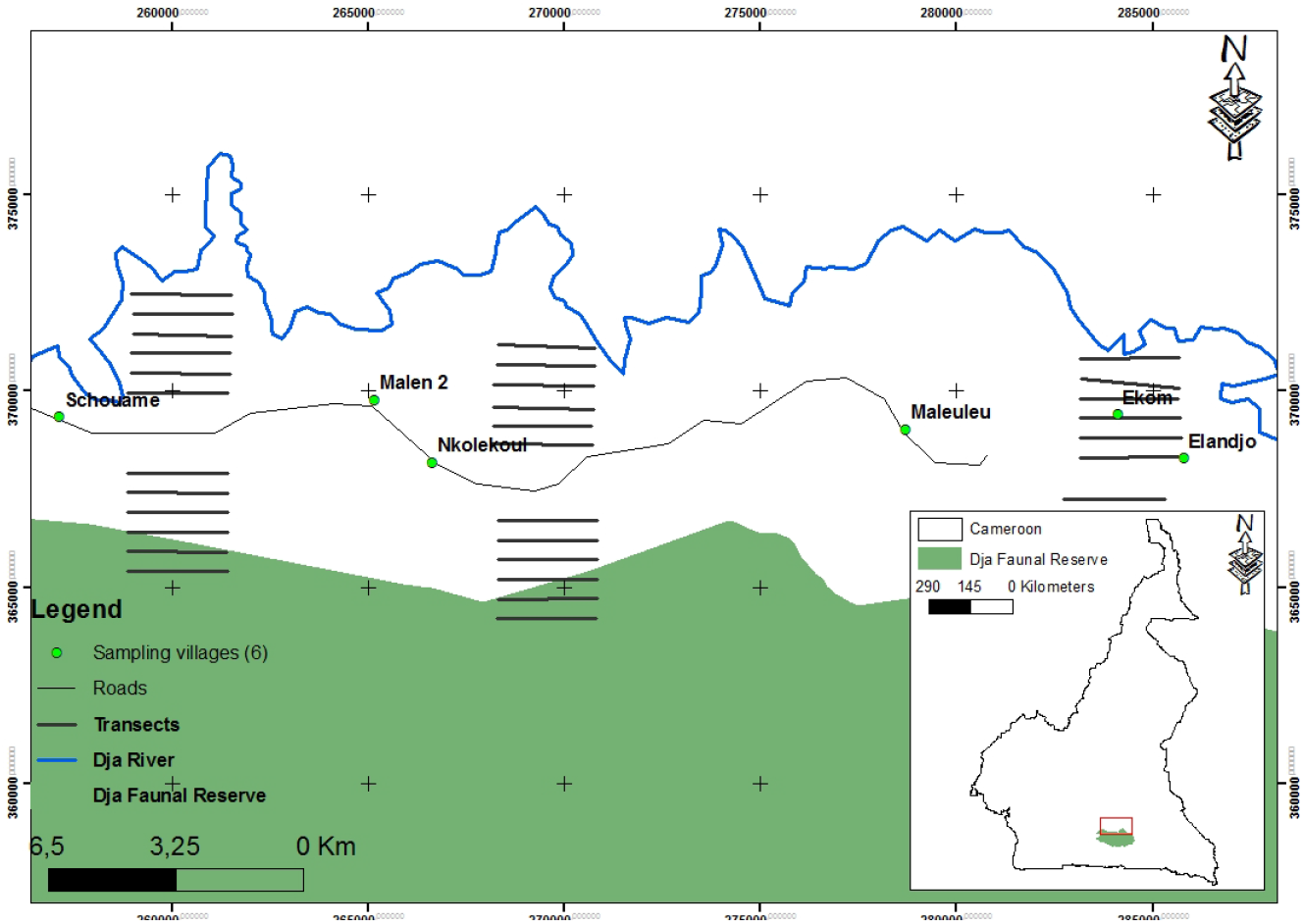
Study area and sampling design presenting transects and sampling villages

### Sampling design

#### Bushmeat offtake

Data on bushmeat offtake were collected in 6 villages (Fig 1) where seven field assistants from the villages were recruited and trained to collect data directly from hunters. Villages were chosen using a stratified random sampling method. The advantage of using locals as assistants is that hunters maintain a better relationship of trust with one of their own. When the hunters left the forest after a hunting party, the assistant was responsible for collecting various information concerning the different species, condition of the carcass (fresh or smoked), weight, time taken for the hunting party, the number of traps set up in the forest by the hunter, and the distance from the village to the location where the animal was killed. For data on hunting parties’ duration, watches were provided to all hunters as well as an information sheet. This was used for recording time of departure and return from hunt while specifying the amount of time devoted to other activities during the trip. The assistant estimated the distance from the village based on known distances between the villages and the Dja River on the one hand and the entrance of the reserve on the other. The boundaries of the reserve are clearly demarcated, therefore allowing hunters to know how far from the village they are. This estimate was based on the hunter’s description of the hunting location. For the entire study duration, the same assistants estimated this distance, thus allowing for comparison over time. A list of local (Badjué) names of all known animals of the area has been established so that all species can be identified and recorded. The hunters were anonymous and just numbers assigned to hunters were recorded by the field assistant. Therefore, based on datasheets, one could not link a specific animal to a hunter; this was done to encourage collaboration of the hunters and thus the collection of data on all animals including protected species. The assistants were regularly visited by a researcher to ensure the quality of data, retrieve the cards, and pay a small fee to each assistant. After each two-month collection round, a distribution of necessities (rice, oil, soap, salt) was organized in all the study villages for all households in order to provide incentives to villagers for a continued participation in the study. Data were collected for eight 2-months periods for three years between December 2017 and December 2020 (Appendix A).

#### Wildlife survey

Three of the 17 villages in the area were selected using a stratified random sampling design in 2018 so as to be distributed throughout the study area. Thus, three sets of 12 transects were opened near the villages in the buffer zone of the DBR (Fig 1). The most distant points of the transects were located at a maximum distance of 4.5 km from the villages and were placed on either side of the trail crossing all villages. Transects were 2.5 km in length and were distributed equally on either side of the trail, resulting in a total length of 30 km per village. The transects were parallel and set 500m apart. The wildlife surveys were carried out in August 2018 and September 2020. Both periods correspond to the rainy season, thus eliminating any bias due to the season and allowing for comparisons between the two years. The surveys were carried out by two teams walking along the transect at an average speed of 1.3km/h. The first team carried out direct observations of the animals and began the inventory between 6.30 a.m. and 7 a.m. Whenever a sighting was made, the name of the species, the distance and the angle of observation, the number of individuals as well as the type of vegetation was recorded. The second team responsible for indirect observations recorded all signs of mammal presence such as dung piles, footprints, hair, and nests. For any animal sign, information such as the species’ name, the type of sign, and the perpendicular distance was noted. In addition to this information, great ape nest data included the name of the plant species used for nest construction, the type of nest, and the age and height of the nest.

#### Data analysis

For each transect, we computed the encounter rate per kilometer (ERK) for human activities by dividing the number of signs by the length of the transect. To analyze the evolution of various bushmeat offtake parameters over time, we calculated for each hunter and for each collection period, the average distance from the village, the number of carcasses, the average hunting duration, and mean body mass indicator (MBMI). Time spent in hunting was calculated by subtracting the time of departure from the time of returning home. In addition, the time allocated to other activities during a period was also deducted. The average hunting duration for each hunter is the total time spent in hunting for a period divided by the number of trips for the same period. We employed the MBMI to investigate changes in the body mass of hunted species (Ávila et al. 2019) over time. This parameter was calculated using the formula

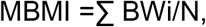

where BWi is the weight of a single individual of species i and N the total number carcasses of all species. The Wilcoxon signed-rank test was used for comparisons of human activities between years and the Kruskall–Wallis test for comparisons of offtake parameters between offtake periods.

The ERK for each species was used to estimate the abundance of wildlife on all transects. An ERK is calculated by dividing the number of signs of a species by the length of the transect (km). We pulled together all types of animal signs to compute the overall abundance for all species as well as that of the different groups (great apes, duikers, small primates, rodents, and pangolins). The Shapiro-Wilk test allowed us to verify if the data were normally distributed. The Wilcoxon signed-rank test was used for comparison of wildlife abundance between the two years. The statistical tests were performed using “Shapiro.test” function (for shapiro test) and wilcox.test function (for Wilcoxon test) of the package “stats” in R 3.4.4 (R Development CoreTeam 2016). Species richness of two years were compared by calculating the diversity estimated for standardized samples using iNEXT Online (Chao et al. 2016), an online version of iNEXT (Interpolation and EXTrapolation) software. We used the asymptotic approach based on interpolation and extrapolation to plot rarefaction curves. These curves, plotted using the number of signs of species (interpolation), are extrapolated to the maximum sampling effort to estimate species diversity. The comparison of wildlife composition between the two years was made using the MRPP, a multivariate permutation analysis testing differences between groups (McCune et al. 2002; Cai 2006; Mota et al. 2010). Analysis were performed with the Sorensen distance measure in PC-ORD 4.0 (McCune and Mefford 1999). In addition, the analysis of indicator species (Dufrêne and Legendre 1997) was carried out in PC-ORD 4.0. This analysis involves the calculation of the indicator value of the species and a test of the statistical significance of this value by permutation tests. A Monte-Carlo test with 0.05 as critical value was used. This test makes it possible to verify whether the occurrence of a species in a given year is significantly different than suggested by a random distribution.

## Results

### Bushmeat offtake

We found a decrease in the number of species targeted by hunters (30 in 2018 to 21 in 2020), total biomass extracted, and number of animals killed. No notable variation was observed in numbers of hunters during the study period (Table 1). Table 2 shows comparisons of different offtake parameters. There was an overall significant reduction in average distance from villages (Kruskall–wallis test, Chi-squared = 58.43; p<0.001). Specifically, we observed a gradual reduction in this distance over time, from 8.07 ± 7.23 km in December 2018 to 7.36 ± 5.97 km in December 2020, with the lowest value recorded in the May–June 2019 period. Comparison of the numbers of carcasses brought back by the hunters showed a significant difference between the different periods (Kruskall–wallis test, Chi-squared = 99.04; p<0.001). Thus, May–June 2019 and December 2019 periods showed an average number of carcasses significantly lower than those of the other periods. There was a drastic reduction in hunting effort. Average hunting duration per hunter of December 2019 (2.95± 2.42 h) was significantly lower than those of July 2018 (5.58 ± 5.27 h) and July 2019 (4.97 ± 4.22 h) (Kruskall–wallis test, Chi-squared = 84.4; p<0.001). Comparison of the average number of traps set by hunters showed a significant overall increase (Kruskall–wallis test, Chi-squared = 120.4; p<0.001). The mean number of hunting trips per hunter did not vary greatly over the study period. The average biomass per hunter was significantly lower in December 2020 (33.53 ± 34.45 kg) compared to the periods of July 2019 (57.93 ± 57.01 kg) and July 2018 (63.00 ± 61.73 kg) (Kruskall–wallis test, Chi-squared = 86.89; p<0.001). Apart from a significantly lower MBMI observed in May 2019 compared to other months, this parameter was held constant throughout the study period.

**Table 1:** General patterns of offtakes: Biomass is the total weight of all species killed by hunters and price the total amount generated by the sale of bushmeat.

| Offtake | Number of species | Number of hunters | Number of carcasses | Biomass (kg) |
| --- | --- | --- | --- | --- |
| 1_ Dec17-Jan18 | 30 | 123 | 592 | 5 991.19 |
| 2 _July18-August 18 | 24 | 116 | 638 | 7 182.06 |
| 3_ December 18- January 19 | 31 | 122 | 609 | 6 901.2 |
| 4_ May 19- June 19 | 30 | 140 | 580 | 5 120.29 |
| 5_ September 19- October 19 | 26 | 140 | 671 | 7 821.61 |
| 6 _December19- January 20 | 23 | 125 | 313 | 3 554.94 |
| 7_ July 20- August 20 | 24 | 137 | 557 | 6 373.09 |
| 8_November 20- December 20 | 21 | 108 | 369 | 4 112.26 |
| **Total** |  |  | **4 329** | **47 016.93** |

**Table 2:** Bushmeat offtake patterns: Numbers in the table are mean number recorded for individual hunters for each offtake period

|  | Offtake1 | Offtake2 | Offtake3 | Offtake4 | Offtake5 | Offtake6 | Offtake7 | Offtake8 |
| --- | --- | --- | --- | --- | --- | --- | --- | --- |
| Distance to village | 8.07 ± 7.23<sup>a</sup> | 7.86 ± 4.77<sup>b</sup> | 6.38 ± 3.9<sup>c</sup> | 4.01 ± 3.42<sup>a,b,c,d,e,f</sup> | 6.95 ± 6.27 | 6.84 ± 5.79<sup>d</sup> | 6.82 ± 5.23<sup>e</sup> | 7,36 ± 5,97<sup>f</sup> |
| Number of carcasses | 30.94 ± 33.95<sup>e</sup> | 50.14 ± 56.94<sup>a,c</sup> | 36.34 ± 44.31<sup>b,d</sup> | 13.72 ± 26.68<sup>c,d,e,f,g,h</sup> | 26,25 ± 33.97<sup>g</sup> | 16.06 ± 22.40<sup>a,b</sup> | 32.81 ± 45.01<sup>f</sup> | 27,38 ± 31,63<sup>h</sup> |
| Hunting duration | 4.89 ± 7.19<sup>a</sup> | 5.58 ± 5.27<sup>c</sup> | 5.21 ± 5.31 | 4.33 ± 4.49 | 4.97 ± 4.22<sup>a,b</sup> | 2.95 ± 2.42<sup>b,c</sup> | 4.15 ± 3.39 | 3,48 ± 2,69 |
| Weight | 49.51 ± 73.51 | 63.00 ± 61.73<sup>b,d</sup> | 58.98 ± 70 | 38.21 ± 42.19<sup>a,b</sup> | 57.93 ± 57.01<sup>a,c</sup> | 33.53 ± 34.45<sup>c,d</sup> | 47.56 ± 45.80 | 38,79 ± 37,4 |
| MBMI | 10.09 ± 5.94 | 11.32 ± 5.27<sup>a</sup> | 10.6 ± 5.77<sup>b</sup> | 8.31 ± 4.46<sup>a,b,c,d,e,f</sup> | 11.02 ± 5.35<sup>c</sup> | 10.71 ± 5.44<sup>d</sup> | 11.16 ± 5.72<sup>e</sup> | 10,32 ± 5,25<sup>f</sup> |
| Number of traps | 41.11 ± 19.91 | 50.65 ± 40.42<sup>b,j</sup> | 46.63 ± 48.35<sup>efg</sup> | 30.44 ± 28.01<sup>a,b,c,d,e</sup> | 64.14 ± 35.01<sup>d,f,h</sup> | 48.82 ± 43.01<sup>a,f,h,i</sup> | 66.24 ± 35.01<sup>e,g,i,j</sup> | 63.32 ± 43.01<sup>c,e</sup> |
| Number of hunting trips/2months | NA | 2.62 ± 2.33 | 2.52 ± 2.02 | 2.78 ± 2.32 | 2.57 ± 1.89 | 2.09 ± 1.39 | 2.28 ± 1.65 | 1.93 ± 1.28 |

<sup>a,b,c,d,e,f,g,h,i,j</sup> numbers on the same line with same letters present significant differences at 5% level, NA: Not Available

### Wildlife abundance

Comparison between years showed a significant difference in overall wildlife abundance, with a higher abundance in 2018 compared to 2020 (W = 831.5; P = 0.039), but differences were minimal (Figure 2). The comparison between groups of species showed no significant difference. However, trends emerged, indicating a decrease in the abundance of large and small primates, rodents, elephants, and pangolins in 2020, while that of duikers increased (Figure 2). Of the 29 species recorded for the two years, five displayed a significant decrease in their abundance between years: bay duiker *(Cephalophus dorsalis)* (p= 0.019), yellow-backed duiker *(Cephalophus sylvicultor)* (p = 0.017), red river hog *(Potamocherus porcus)* (p < 0.001), flat-headed kusimance *(Crossarchus platycephalus)* (p = 0.007), and a squirrel (Funisciurus sp) (p = 0.021) (Appendix A).

**Figure 2:**
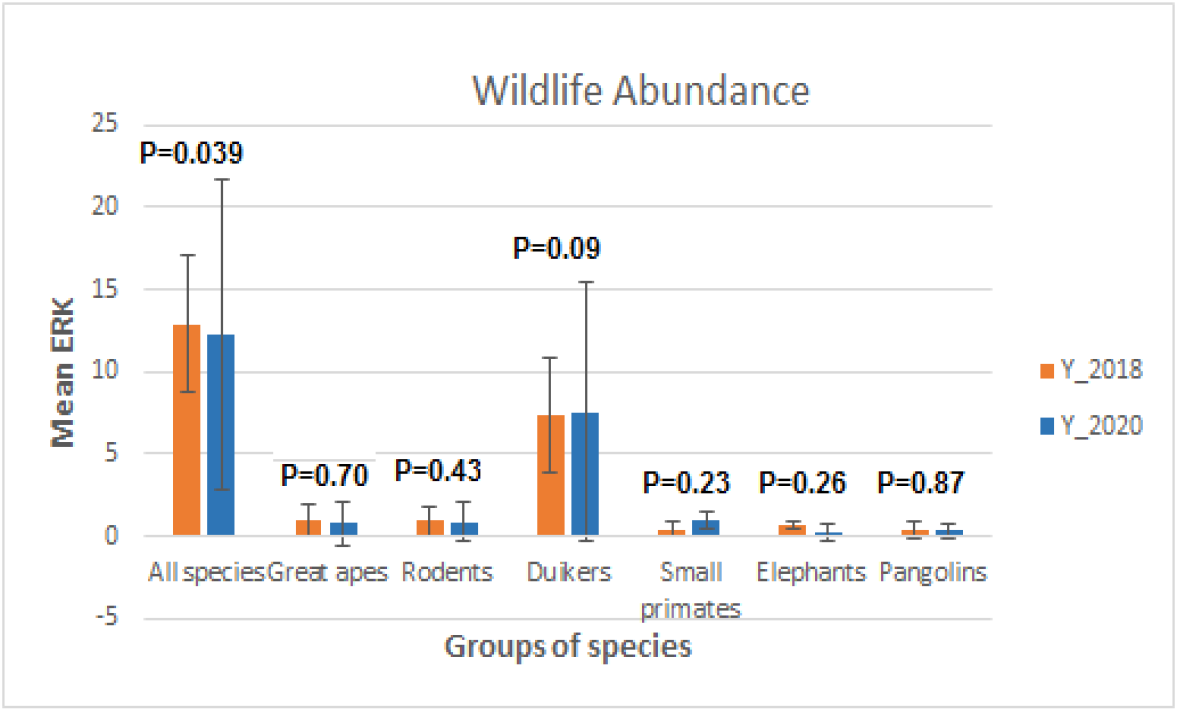
Comparison of wildlife abundance between years. ERK is encounter rate per kilometer.

### Wildlife species richness

We recorded 26 and 23 species in 2018 and 2020, respectively (Appendix A). The species richness of the area did not change significantly over the two years of the study. Rarefaction and extrapolation curves were fitted based on wildlife sightings and did not show any significant differences (p > 0.05) in species richness between study years as the 95% confidence intervals overlap (Fig. 3). The confidence intervals converge to 23–29 and 21–31 for years 2018 and 2020, respectively.

**Figure 3:**
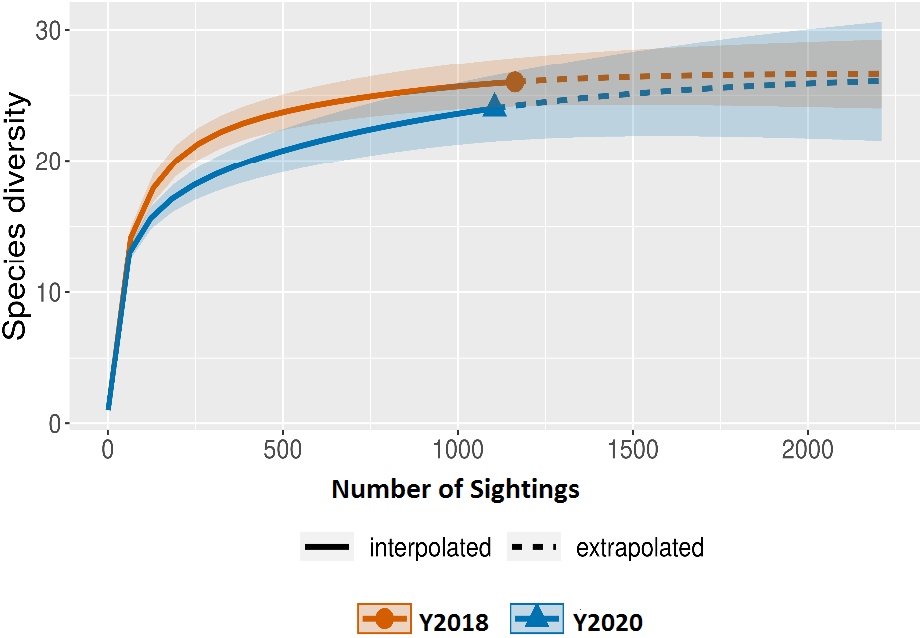
Rarefaction and extrapolation curves comparing wildlife species richness. The 95% confidence intervals (CIs) converge to 23–29 and 21–31 respectively for Years 2018 and 2020. There is no significant difference (p > 0.05) between years as the CIs overlap.

### Wildlife composition

Mammal community composition for both study years overlapped greatly: a total of 29 species was recorded, and 20 occurred in both years. Compositional differences were statistically significant, as shown by MRPP results (T = −8.39, p< 0.001). However, the indicator species analysis showed that, unlike the species recorded in 2020 which did not present any significant value, flat-headed kusimanse *(Crossarchus platycephalus)* (p = 0.01), water chevrotain *(Hyemoschus aquaticus)* (p = 0.03), and red river hog *(Potamochoerus porcus)* (p = 0.005) were the indicator species of the year 2018 (Appendix A).

## Discussion

Evaluating the effectiveness of conservation actions requires information derived from different types of data to predict hunting sustainability (Van Vliet and Nasi 2008). Monitoring the temporal trends of both the bushmeat offtake and the wildlife community may help detect suitable indicators for a rapid assessment of the effectiveness of conservation actions. Our study presents some limitations mainly linked to the one-off surveys on transects which do not always guarantee that all species present in the environment have been listed. In addition, the transect method seems less suitable for the survey of some species. However, the same methodology was used for the two inventories of our study, making our results comparable.

Results show a considerable drop in the number of species brought back to the village by hunters. This decrease may be due to a better awareness of hunting restrictions according to the Cameroonian law. This explanation is plausible since the results of wildlife surveys show that species that were no longer hunted persist in the area and the number of hunters remained unchanged during the different periods. This awareness may result from the activities of several conservation organizations in the area. In addition, the number of hunted species is relatively low compared to the number reported for neighboring areas (Ávila et al. 2019). These results corroborate our hypothesis of the reduction in the number of species hunted. The results also show a reduction, over time, of the hunting distance and the time spent hunting. This may be explained by the increased involvement of local people in agricultural activities, which limits the amount of time dedicated to other activities. At the end of our study, the time spent in hunting activity was relatively low compared to that of other villages in the region where hunters undertook day-long trips, spending an average of 4.6h per week hunting (Epanda et al. 2005). This reduction in hunting effort supports our assumptions. The reduction in time and distance logically leads to a decrease in the average biomass brought back by the hunters, as presented in our results, because hunters reduced their hunting effort. This reduction in biomass contrasts with results obtained in other areas in Africa where the tendency was rather the increase in hunting (Fa and Yuste 2001). Our results show that, except for the period of May–June 2019, the average MBMI in the study villages did not drop over time. The low value of MBMI in this period could be explained by the fact that the government of Cameroon suspended the sale of cartridges during this period, thus reducing hunting with firearms. Results therefore indicate that our study area is not overhunted, and there is no preponderance of smaller-bodied game species (Fa et al. 2015; Ingram et al. 2015) compared to other villages in the region (Ávila et al. 2019). Hence, body masses of carcasses brought to the villages every day were similar throughout the study period, suggesting that prey populations were not depleted (Avilla et al 2019). Trends in MBMI are consistent with our hypothesis of negligible change in that value. A slight but significant increase was observed in mean number of traps set. This could be explained by the fact that the traps are set only once in the forest and can be used for more than a year. Therefore, even if the hunters no longer visit them all because of the reduction in time and distance they cover, they still consider that they have the same number of traps in the forest and sometimes hunters can set more traps near villages. Our results suggest that the behavior of hunters was modified throughout the study period given the decrease in harvested biomass and the decrease in the hunting distance and time spent in hunting activity.

Overall wildlife abundance was lower in 2020 compared to 2018, but the difference was only 5% higher in 2018. However, no differences in the abundance of the different groups were observed. Regarding primates, there was no differences in great ape and small primate abundance across the years. No significant difference in the abundance of rodents was observed between the two years. Specifically, five species have shown a reduction in their abundance over the years: three medium-sized species (bay duiker, yellow-backed duiker, and red river hog) and two small mammals (flat-headed kusimance and squirrels). The reduction in the abundance of medium-sized mammals may be due to hunting pressure in the area. In fact, hunters use firearms for hunting and therefore select species which have a large body mass because the main objective of hunting is meat. So, the abundance of these species which are sensitive to hunting is likely to decrease. Moreover, species such as the yellow-backed duiker are rare and even extinct in areas close to the Dja Faunal Reserve (Bobo et al. 2014). The overall decline in abundance reflects the general situation of tropical forests (Beaudrot et al. 2016a). For example, in some areas in Amazonia, declines in some wildlife species occupancy and abundance were noted (Ahumada et al. 2011; Beaudrot et al. 2016b); this was also the case for carnivores in some parts of Asian tropical forests (Jenks et al. 2011). Nevertheless, the observed decrease is minimal compared to the situation in the nearby area where a decrease in abundance of more than 20 percent and local extinction of some species have been observed over a 14 year-period (Tagg et al. 2020). Our results show that great ape populations remained stable, but this could be due to the small interval between the surveys because several studies in wildlife monitoring in tropical forests have been conducted in the minimum periods of 3 to 8 years (Jenks et al. 2011; Beaudrot et al. 2016a). Small primates and rodent abundance also remained stable throughout the years. In addition to being small and therefore less interesting for hunters (Ráez-Luna 1995), small primates have large populations and a high reproduction rate and exhibit population-level density compensation in response to extirpation, making them more resilient to hunting disturbances (Peres 1990; Rosin and Swamy 2013). Rodents as small animals provide less incentive for hunting in an area where the goal of hunting is high bushmeat offtake for commercial goals. Moreover, because of their ability to reproduce quickly, rodent populations can thrive in hunted areas (Berryman 1992; E. Bennett and J. Robinson 2000; Abernethy et al. 2013). Hunting pressure in protected areas is lower than in the buffer zone where the strong hunting pressure is associated with a higher percentage of large rodents offtake (Ziegler et al. 2016). These observations are not completely in line with our expectations that the overall abundance would remain constant over the years given the slight decrease in abundance. Despite the decline in overall wildlife abundance over the years, individual-based rarefaction curves suggest that reduction of wildlife abundance did not significantly affect wildlife species richness. Constant values of species richness are consistent with the trends previously observed in the area, showing that it takes a relatively long period to observe a change in species richness (Tagg et al. 2020). However, such change may result from an increase in hunting pressure, which we have demonstrated in this study to have reduced (Appendix B). This result suggests that no local extinction occurred despite the decrease in wildlife abundance – and further suggests that if sustained over time, conservation efforts should help to increase wildlife metrics. We also found a slight but statistically significant change in wildlife composition of the area over the years, thus supporting previous studies showing a change in wildlife composition in the northern edge of DBR over the time (Muchaal and Ngandjui 1999; Tagg et al. 2020). Such compositional changes corroborate earlier studies that underline a selection in hunted species (Abernethy et al. 2013; Ripple et al. 2016) in which some species are less resilient than others. For example, larger-bodied species with slower reproductive rates (Arnhem et al. 2008; Stokes et al. 2010; Linder and Oates 2011). All functional diversity indices remained constant throughout the study period. This is mainly due to the fact that these indices, although included in the parameters related to species ecology, are also based on factors such as species richness and wildlife community composition and abundance and therefore follow the same trends as the former. Given that recorded offtake data at the same period show that all the species were still present in the area, we can conclude that these species are not depleted but increasing hunting pressures may lead to extinctions (Ripple et al., 2016). These results of species richness, wildlife composition, and functional diversity partially support our hypotheses which predicted no change for all these parameters.

Our results suggest that the monitoring of parameters related to bushmeat offtake and hunting exhibits more rapid changes than those related to wildlife abundance for approximately the same time interval. Indeed, the number of species hunted, the number of carcasses and the associated biomass presented a significant decrease over time. In addition, hunting effort also presented a gradual reduction during the study period. These observed variations may be due to the fact that these different indices are mainly related to hunters’ behavior that is likely to change in a relatively short period of time in a context where alternatives to hunting activity are available. In the area, these alternatives mainly consist of support in cocoa production and the supply of fishing equipment that appear as supplements to activities that rural people already carry out. This is supported by the fact that in the northern outskirts of the Dja Faunal Reserve, it has been shown that hunting pressure is not determined by the distance or accessibility of the site but rather by hunters behavior (Tagg et al. 2015). Therefore, these alternative activities are likely to change their hunting behavior more quickly (Ferraro 2001; Ferraro and Kiss 2002; Winkler 2011). Our model corroborates that of Winkler (2011) and Salafsky (2011) who showed that focusing on the social aspect in conservation projects produces more significant changes than simply taking into account the development of means to avoid wildlife extraction from forests. Unlike bushmeat harvest trends, we observed little variation with regard to wildlife abundance in the forest as well as the various indices of functional diversity. This contrast in the level of variation of various parameters over time implies that bushmeat offtake may be the best indicator for short-term evaluation of conservation projects. These results suggest that if a low level of bushmeat extraction is maintained over time, the wildlife population will increase. This inverse relationship between hunting and future change in wildlife community structure is corroborated by the fact that gradual increase in bushmeat extraction leads over the years to a decrease in functional diversity (Ávila et al. 2019; Tagg et al. 2020). In tropical forests, raising awareness of the importance of conserving great apes has led to a reduction in hunting pressure and as a result to an increase in their populations (Sandbrook and Roe 2010; Tagg et al. 2011). In addition, studies show that in sites where hunting pressure is low, there is a relative increase over time in wildlife abundance (Effiom et al. 2013; Fa et al. 2015; Tagg et al. 2015). This demonstrates the value of monitoring bushmeat/hunting as metrics that will easily fluctuate in the short term because they are more quickly altered as a result of community-level conservation actions; with this sustained change over time, we would then confidently expect to see a positive change of wildlife metrics.

## Conclusion

Our study emphasizes the importance of using data from hunting for the short-term evaluation of conservation actions. It was based on the monitoring of bushmeat offtake data and wildlife survey in order to determine the index which exhibits more rapid variations over time. The monitoring of bushmeat harvesting dynamics over the three years has shown a significant overall reduction in the biomass extracted, time devoted to hunting and distance traveled by hunters. However, no significant variation was observed with regard to the body mass of hunted species and the number of traps set. We found an overall but slight reduction in wildlife abundance as well as a change in wildlife community composition over the two years of study. However, wildlife species richness and functional diversity remained unchanged. Our results reveal that bushmeat offtake parameters are more likely to vary between short periods compared to parameters related to the living wildlife community. Our results underscore the need to focus more attention on species extraction patterns and hunters’ behaviors when evaluating the effectiveness of wildlife conservation actions undertaken over short-term periods in areas subject to hunting. Ecological monitoring of living wildlife may require more time to effectively detect change and guide the evaluation of such projects. In a context where the major concern of conservationists is the reduction of hunting, the successful evaluation of the effectiveness of conservation actions inevitably depends on monitoring of factors linked to this activity. This study contributes to the identification of suitable indicators for the evaluation of conservation actions.

## Acknowledgements

We thank the Ministry of Forestry and Wildlife and the Ministry of Scientific Research and Innovation for authorizing this research. We also thank the Conservator of Dja Biosphere Reserve for collaboration during this work. Data collection would not have been completed without the ultimate contribution of many research assistants, namely: Christian Nguenga, Germain Kameni, and Constantin Deutchoua. We thank all the PGS staff for their administrative and logistic support of the research: Bernadette Bayimbe, God Love Mbunwe, and Honorine Nyanda, speciel thank to Luc Tedonzong. We also acknowledge the undeniable contribution of local guides to the implementation of this research project, namely: Serges Nguemo, Sakanla, Biance Moyetina, Michel Olé, and many others.

**Appendix A:** Indicator species analysis and ERK comparisons of all species between 2018 and 2020.

|  | Encounter rate per kilometer (ERK) |  |  |  | Indicator species analysis |  |  |  |
| --- | --- | --- | --- | --- | --- | --- | --- | --- |
| Species | 2018 | 2020 | Wilcoxon test | P | Modal year | Indicator Value | Standard Deviation | P-value |
| *Aonyx sp* | 0.02 ± 0.07 | 0 ± 0 | 630 | 0.333 | 2018 | 2.8 | 0.09 | 1.0000 |
| *Atherurus africanus africanus* | 0.64 ± 0.64 | 0.81 ± 1.17 | 608.5 | 0.96 | 2020 | 38.8 | 4.78 | 0.4370 |
| *Bdeogale nigripes* | 0.03 ± 0.09 | 0 ± 0 | 647.5 | 0.16 | 2018 | 5.6 | 2.09 | 0.5000 |
| *Cephalophus callipygus* | 1.16 ± 1.02 | 1.6 ± 1.79 | 588 | 0.77 | 2020 | 37.0 | 4.45 | 0.8350 |
| *Cephalophus dorsalis* | 2.96 ± 1.92 | 2.88 ± 3.47 | 795.5 | 0.031* | 2018 | 51.2 | 3.90 | 0.4990 |
| *Cephalophus monticola* | 1.78 ± 1.48 | 1.70 ± 2.04 | 726.5 | 0.179 | 2018 | 50.0 | 4.18 | 0.4540 |
| *Cephalophus nigrifrons* | 0.21 ± 0.33 | 0.36 ± 0.56 | 544 | 0.367 | 2020 | 28.4 | 4.46 | 0.2590 |
| *Cephalophus sylvicultor* | 0.99 ± 0.64 | 0.91 ± 1.26 | 795 | 0.029* | 2018 | 49.1 | 4.27 | 0.0810 |
| *Cercopithecus cephus cephus* | 0.05 ± 0.12 | 0.28 ± 0.89 | 587 | 0.613 | 2020 | 11.6 | 3.76 | 0.4470 |
| *Cercopithecus nictitans nictitans* | 0.17 ± 0.37 | 0.15 ± 0.21 | 610.5 | 0.982 | 2018 | 15.4 | 4.00 | 0.5900 |
| *Cercopithecus pogonias* | 0.09 ± 0.19 | 0.08 ± 0.25 | 569.5 | 0.387 | 2018 | 10.4 | 3.39 | 0.5720 |
| *Cricetomys emini* | 0.14 ± 0.29 | 0.07 ± 0.18 | 669 | 0.333 | 2018 | 14.8 | 3.63 | 0.3380 |
| *Crossarchus platycephalus* | 0.14 ± 0.27 | 0.01 ± 0.06 | 753 | 0.006* | 2018 | 23.1 | 3.28 | 0.0100* |
| *Gorilla gorilla gorilla* | 0.15 ± 0.43 | 0.04 ± 0.16 | 682.5 | 0.179 | 2018 | 14.9 | 3.57 | 0.2400 |
| *Herpestes naso* | 0.09 ± 0.19 | 0.02 ± 0.09 | 701 | 0.074 | 2018 | 15.6 | 3.40 | 0.1410 |
| *Hyemoschus aquaticus* | 0.01 ± 0.02 | 0 ± 0 | 612.5 | 1 | 2018 | 17.7 | 3.13 | 0.0380* |
| *Cercocebus sp* | 0.02 ± 0.09 | 0 ± 0 | 647.5 | 0.16 | 2018 | 5.6 | 2.09 | 0.5000 |
| *Loxodonta africana cyclotis* | 0.07 ± 0.18 | 0.25 ± 0.54 | 547 | 0.26 | 2020 | 17.5 | 3.72 | 0.1710 |
| *Manis gigantea* | 0.18 ± 0.31 | 0.13 ± 0.24 | 655.5 | 0.528 | 2018 | 18.1 | 3.91 | 0.5170 |
| *Manis tricuspis* | 0.2 ± 0.34 | 0.21 ± 0.33 | 602 | 0.889 | 2020 | 18.1 | 4.31 | 0.9470 |
| *Cephalophus sp* | 0 ± 0 | 0.02 ± 0.07 | 595 | 0.331 | 2020 | 2.8 | 0.09 | 1.0000 |
| *Funisciurus sp* | 0 ± 0 | 0.03 ± 0.09 | 577.5 | 0.16 | 2020 | 5.6 | 2.09 | 0.5010 |
| *Nandinia binotata* | 0.023 ± 0.09 | 0.02 ± 0.09 | 612.5 | 1 | 2018 | 2.8 | 2.64 | 1.0000 |
| *Neotragus pygmaeus* | 0 ± 0 | 0.02 ± 0.09 | 577.5 | 0.16 | 2020 | 5.6 | 2.09 | 0.5170 |
| *Pan troglodytes troglodytes* | 0.48 ± 0.68 | 0.73 ± 1.33 | 567 | 0.572 | 2020 | 33.4 | 5.12 | 0.3760 |
| *Panthera pardus* | 0.02 ± 0.07 | 0 ± 0 | 630 | 0.333 | 2018 | 2.8 | 0.09 | 1.0000 |
| *Potamochoerus porcus* | 2.94 ± 1.34 | 1.97 ± 1.5 | 879 | 0.001* | 2018 | 60.0 | 2.89 | 0.0050* |
| *Funisciurus sp* | 0.06 ± 0.14 | 0 ± 0 | 700 | 0.021* | 2018 | 13.9 | 2.58 | 0.0650 |
| *Tragelaphus spekei* | 0.33 ± 0.47 | 0.29 ± 0.58 | 683 | 0.349 | 2018 | 25.0 | 3.5 | 0.5700 |

**Appendix B:** Human activities abundance comparison between the two years of study

| **Human activities** | **2018** | **2020** | **W** | **P** |
| --- | --- | --- | --- | --- |
| All signs | 1.39 ± 0.94 | 1.58 ± 0.93 | 566.5 | 0.35 |
| Hunting signs | 0.51 ± 0.5 | 0.76 ± 0.62 | 962.0 | 0.087 |
| Other human signs | 1.01 ± 0.9 | 1.04 ± 0.7 | 655.5 | 0.608 |

